# Transcranial direct current stimulation alters functional network structure in humans

**DOI:** 10.1101/296657

**Authors:** M. Ruttorf, S. Kristensen, L.R. Schad, J. Almeida

## Abstract

Transcranial direct current stimulation (tDCS) is routinely used in basic and clinical research, but its efficacy has been challenged on a methodological and statistical basis recently. The arguments against tDCS derive from insufficient understanding of how this technique interacts with brain processes physiologically. Because of its potential as a central tool in neuroscience, it is important to clarify whether and how tDCS affects neuronal activity. Here, we investigate influences of offline tDCS on network architecture measured by functional magnetic resonance imaging. Our results reveal a tDCS-induced reorganisation of a functionally-defined network that is dependent on whether we are exciting or inhibiting a node within this network, confirming in a functioning brain, and in a bias free and independent fashion that tDCS influences neuronal activity. Moreover, our results suggest that network-specific connectivity has an important role in defining the effects of tDCS and the relationship between brain states and behaviour.

## Introduction

Transcranial direct current stimulation1,2,3,4,5 (tDCS) has been widely used in the neurosciences6,7,8,9 for decades. This is so because interfering techniques like tDCS that are assumed to directly modulate neuronal activity are extremely promising for both basic and applied research as they allow for addressing research questions on the causal relationships between brain states and behaviour10,11,12. However, the efficacy of tDCS has been put into question recently13,14,15,16,17 on a methodological and statistical basis. It is thus central to have a closer look at the effects of tDCS on brain activity. Previously, we provided evidence that offline tDCS locally affects neuronal responses in accordance with stimulation polarity (i.e., inhibition or excitation) as measured by functional magnetic resonance imaging (fMRI)5. Nevertheless, the global effect of tDCS on functional brain networks in humans is still not well understood18,19, but is central for a better and more informative understanding of the mechanisms of tDCS. Based on these previous findings and on the detailed work on living macaques by Krause et al. (2017)20, here we decided to investigate, in humans, the outcome of tDCS on the underlying functional architecture of the brain as measured by fMRI. There are certain key methodological issues related to the effect of tDCS in the brain that are currently unsolved21,13,12. These include understanding the technique’s (i) functional focality, i.e. is tDCS limited to local effects on the stimulated area, or do the effects also transfer more globally to the network level as pointed out by Krause et al (2017)20; (ii) specificity of stimulation, i.e. is tDCS-induced interference dependent on general processes such as the spatially wide expansion of the electrical field22, or is it dependent on more neuronally-specified processes such as functional connectivity between regions; or (iii) modulatory effects, i.e. how does tDCS modulate functional connectivity between brain regions. Up to now, there are only two studies evaluating the effect of tDCS on the structure of underlying functional brain networks in depth by means of graph theory: one uses tDCS in combination with resting-state fMRI23, and the other combines tDCS with electroencephalography24. Importantly, none of these examined topology changes in functional brain networks in detail. For this reason and because cognitive functions rely on the processes happening within networks of functionally-connected brain regions rather than on local and isolated areas, we look at how tDCS affects neuronal organisation using a task-based fMRI experiment in combination with offline tDCS. We did so because: (i) task performance enhances neuronal activity resulting in functional connectivity between relevant brain areas being more reliable in terms of graph theory metrics25; (ii) tDCS preferentially modulates active neuronal networks, when compared to inactive networks sharing the same anatomical space (activity-selectivity approach)26; and (iii) offline tDCS allows us to map the spatio-temporal patterns of functional reorganisation at the systems level27.

### Experimental Layout

We combined tDCS with a task-based paradigm in fMRI using a repeated measures design (see Methods for more details). We asked a group of ten individuals to participate in four experimental sessions, each one separated by at least one week. In the first session, participants went through the fMRI experiment only – as control session – whereas in the second to fourth sessions participants were first subject to tDCS stimulation outside the MR scanner that was immediately followed by the fMRI paradigm. The paradigm consisted on passively watching pictures of tools, animals, faces and places. The tDCS sessions consisted of anodal (typically thought to increase neuronal excitability) or cathodal (typically thought to decrease neuronal excitability)28,19 stimulation to either the left Inferior Parietal Lobule (IPL) or the right Superior Temporal Sulcus (STS). This resulted in four experimental within-participant groups: anodal stimulation on IPL (AnoIPL), cathodal stimulation on IPL (CatIPL), cathodal stimulation on STS (CatSTS) and control (Ctrl). We chose the left IPL and right STS as target areas because they are highly accessible to the tDCS stimulation technique. Moreover, IPL is known to respond more to images of tools than images of stimuli from other categories29, whereas STS does not30. This is important because by using STS we obtained a tDCS “sham” group to compare tDCS to IPL with – additionally to the control group that serves as ground truth without stimulation. Contrary to classical sham procedures, here participants receive active stimulation to an alternative location to counter doubts which arose recently31,13 concerning the ability to distinguish classical sham from active stimulation.

We decided to concentrate on brain areas that are dedicated to the processing of tool items (i.e., the tool network5,32,33), which left IPL is an exemplary constituent, because effects of tDCS depend on the cognitive/neural processing participants are engaged in – i.e., because this network would be actively processing the tool stimuli presented in our experiment, we could better test the effects of tDCS over this global network. We selected 18 regions of interest (ROIs) that have been associated with tool processing29,34,35. The location of the ROIs can be seen in Figure 1 using BrainNet Viewer software36 (Version 1.53) as red spheres placed on the ICBM-152 template37. The location corresponds to the ROIs’ centre coordinates listed in Table 1. Brain networks demonstrate hierarchical modularity (or multi-scale modularity) - i.e. each module contains a set of sub-modules that contains a set of sub-sub-modules, etc38. Object recognition – and thereby the tool network as well – is organised in a modular way comparable to colour vision which is shown to be automatic, effortless and informationally encapsulated39. Thus, we treated the tool network as a modular network with a subset of highly functional-connected nodes. Keeping this in mind, we are able to test whether tDCS can induce reorganisation over a functional network in the brain, and specifically here over the tool network, beyond the known local effects over the stimulated area5.

**Table 1.** Overview of brain regions in analysed functional network.

| Lobe | Hemisphere | Structure | Brodmann<br>area [BA] | Label | Talairach Coordinates |  |  |
| --- | --- | --- | --- | --- | --- | --- | --- |
|  |  |  |  |  | X | Y | Z |
| ROIs of Tool Network |  |  |  |  |  |  |  |
| temporal | left | posterior middle<br>temporal gyrus | BA 37 | pMTG | -42 | -62 | -7 |
| temporal | left | middle fusiform<br>gyrus | BA 37 | MFG | -24 | -48 | -8 |
| occipital | left | extrastriate<br>visual cortex | BA 19 | V3aV7 | -23 | -80 | 24 |
| occipital | right | extrastriate<br>visual cortex | BA 19 | V3aV7 | 25 | -78 | 27 |
| parietal | left | anterior angular<br>gyrus | BA 39 | PGa | -41 | -63 | 33 |
| occipital | left | posterior<br>angular gyrus | BA 39 | PGp | -40 | -73 | 24 |
| parietal | left | superior parietal<br>lobe (anterior<br>parts) | BA 7 | BA7a | -18 | -64 | 49 |
|  | right |  |  |  | 18 | -65 | 50 |
| parietal | left | superior parietal<br>lobe (posterior<br>parts) | BA 7 | BA7p | -10 | -76 | 40 |
|  | right |  |  |  | 13 | -76 | 44 |
| parietal | left | lateral superior<br>parietal lobe | BA 7 | 7PC | -31 | -55 | 50 |
| parietal | right | lateral superior<br>parietal lobe | BA 7 | 7PC | 27 | -54 | 49 |
| parietal | left | supramarginal<br>gyrus | BA 40 | PF | -53 | -41 | 31 |
| temporal | left | supramarginal<br>gyrus | BA 40 | PFcm | -45 | -39 | 21 |
| parietal | left | supramarginal<br>gyrus | BA 40 | PFm | -48 | -54 | 34 |
| parietal | left | supramarginal<br>gyrus | BA 40 | PFop | -53 | -28 | 24 |
| parietal | left | supramarginal<br>gyrus | BA 40 | PFt | -48 | -29 | 33 |
| parietal | left | intraparietal<br>sulcus | BA 7/40 | hIP1 | -34 | -52 | 34 |
| parietal | left | intraparietal<br>sulcus | BA 7/40 | hIP2 | -42 | -44 | 37 |
| parietal | left | superior parietal<br>lobe | BA 7/40 | hIP3 | -29 | -54 | 38 |
| Stimulation Sites |  |  |  |  |  |  |  |
| parietal | left | inferior parietal<br>lobe | BA 40 | IPL | -38 | -37 | 36 |
| temporal | right | superior<br>temporal sulcus | BA 37 | STS | 44 | -45 | 6 |

Here, we list the names of the brain areas, labels used in the text and centre coordinates (x,y,z) in Talairach space of the regions of interests (ROI) of the tool network and the stimulation sites. The anterior and posterior parts of the superior parietal lobe are bilateral ROIs.

**Figure 1:**
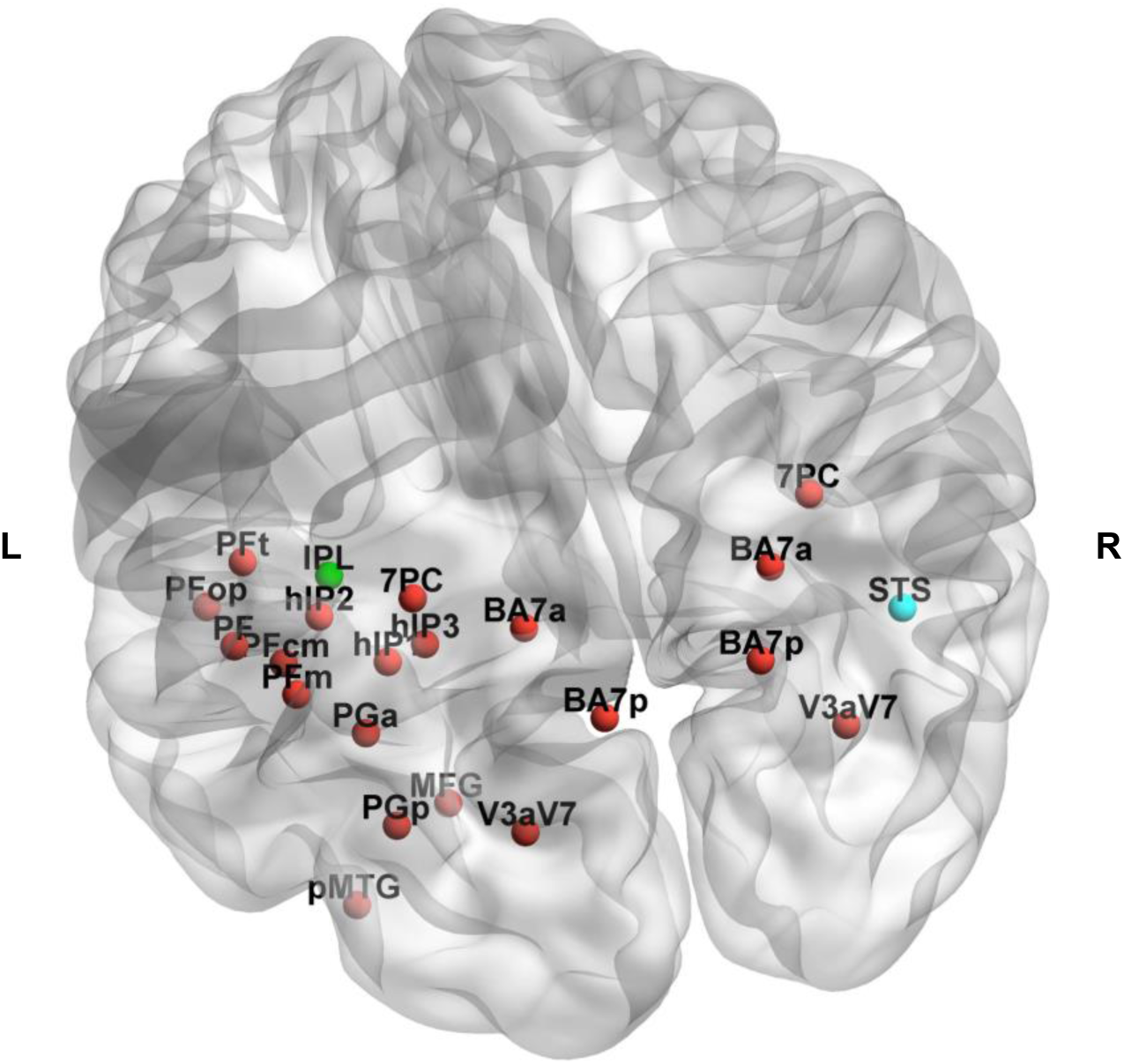
Location of the regions of interest analysed. Coloured in red are the regions of interest (ROIs) within the tool functional network according to centre coordinates and labels given in Table 1. The location of the stimulation sites is shown either in green (Inferior Parietal Lobule – IPL) or in blue (Superior Temporal Sulcus – STS). L/R denotes the left and right hemisphere, respectively. A video with 360° view of the location of the ROIs is available as Supplementary Information.

### Graph Theory Analysis

A graph is mathematical description of a network consisting of nodes *N* (here: the ROIs selected) and edges *k* (here: functional “links” between pairs of ROIs). Below, we refer to graphs explicitly because this does not make any assumptions on the nature of the edges but rather emphasises the aspect of mathematical modelling because “network” generally refers to real-world connected systems40. We analysed weighted undirected graphs averaged per group (see Methods for details of graph construction) using Brain Connectivity Toolbox41 implemented in MATLAB R2013a (The MathWorks Inc., Natick, MA, USA). Because we were interested in changes in underlying network architecture in the brain between experimental groups we looked at topological graph metrics as community structure and participation coefficients primarily. After graph construction, we checked for *N*,*k*-dependence (see Methods). The number of nodes stays constant (*N* = 18) in all experimental groups, the number of edges is almost equal between groups (Δ*k* = ±2, *k*_*max*_ = 153). Using a repeated measures design, we were only interested in changes between experimental groups. So, we kept the resulting graphs while considering the gain or loss of an edge as an effect of the stimulation (tDCS).

### Community Structure

Community structure has been identified as a sensitive marker for organisation in brain networks42. Community structure analysis detects the groups of regions more densely connected between them than expected by chance. The resulting group-level community structure was visualised by assigning a different colour to each community (see Figure 2). This was then displayed by overlaying spheres coloured by community affiliation on the ICBM-152 template as done in Figure 1.

**Figure 2:**
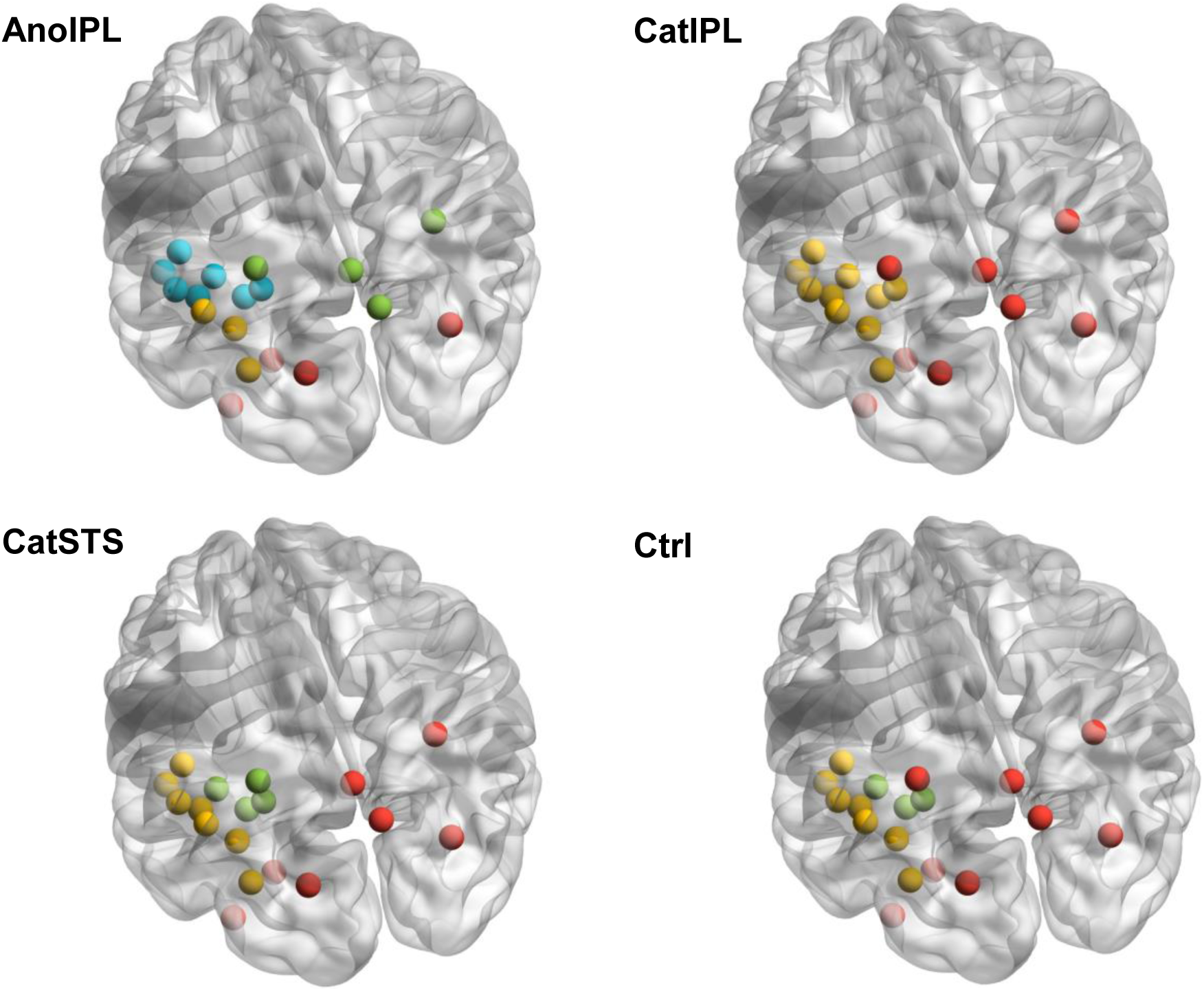
Community structure of the tool network. Within the four experimental groups (AnoIPL, CatIPL, CatSTS and Ctrl), resulting community structures of the tool network are shown. Colours denote different communities; red indicates community I, yellow community II, green community III and blue community IV. Angle of vision kept as in Figure 1. Location of the spheres visualised according to centre coordinates given in Table 1.

The values of modularity *Q* corresponding to the community structures shown in Figure 2 are almost identical (ΔQ = ±0.02). There are three communities in the Ctrl and CatSTS experimental groups, two in CatIPL experimental group and four in AnoIPL experimental group. The communities in Ctrl and CatSTS experimental groups differ minimally from each other. One node changed community assignment (from community III to community I). In AnoIPL experimental group, the community structure intensifies to four whereas in CatIPL experimental group the community structure relaxes to two. We controlled for possible limitations43 relevant to our experimental layout: the results shown in Figure 2 are neither subject to resolution limit of the objective function44 nor dependent on the method used to average the correlation coefficients (see Methods for more details). Furthermore, we overlaid the community structure for each experimental group on their averaged weighted temporal correlation matrix before converting to absolute values to verify that negative edge weights are sparser within and denser between communities found45. Likewise, we overlaid the community structure for each experimental group on their distance matrix (see Methods) to re-examine that distances within communities are smaller than between communities as shown in Figure 3. We show that the number of communities changed depending on stimulation site and polarity of tDCS. While there is almost no difference in community affiliation when stimulating STS which does not belong to the tool network, there are clear polarity-dependent effects when stimulating IPL.

**Figure 3:**
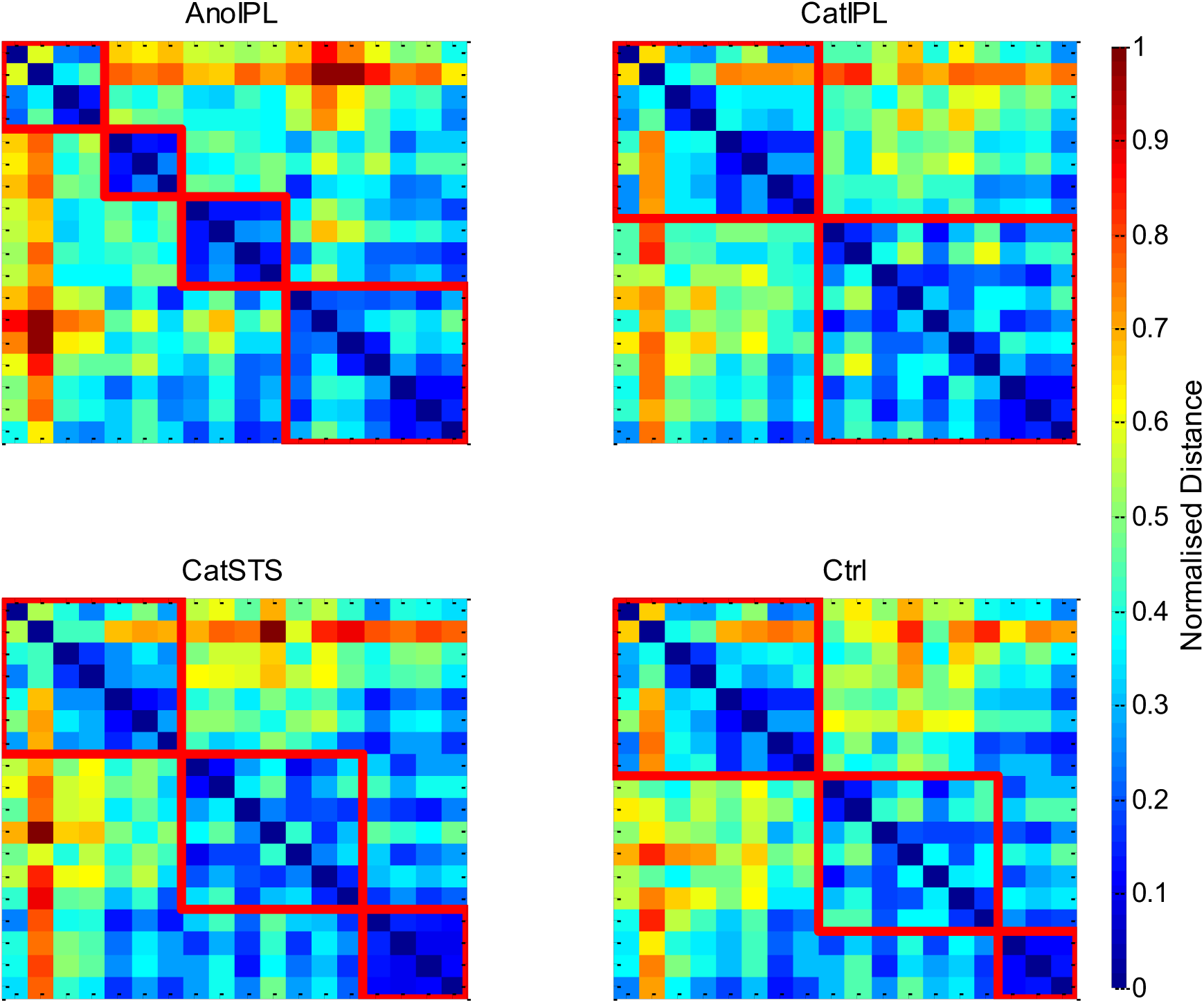
Plots of distance matrices with community structure on top. For the four experimental groups (AnoIPL, CatIPL, CatSTS and Ctrl), normalised distance matrices grouped by communities are shown. The borders of the communities are marked by thick red lines. The colour bar indicates the normalised distance between nodes. The distance is less within communities than between communities throughout experimental groups in all communities found.

### Participation Coefficient

While the within-module degree *z* score defines the role of a node in its own community, the participation coefficient *P* is a feature of each node’s connectivity relative to the community structure of the entire network46. Nodes with a low value of *P* share connections with other members of the same community, whereas those with a high *P* value serve as connectors between communities. In Figure 4, the *P* values for the four experimental groups are plotted in the *P-z* parameter plane (see Methods for details).

**Figure 4:**
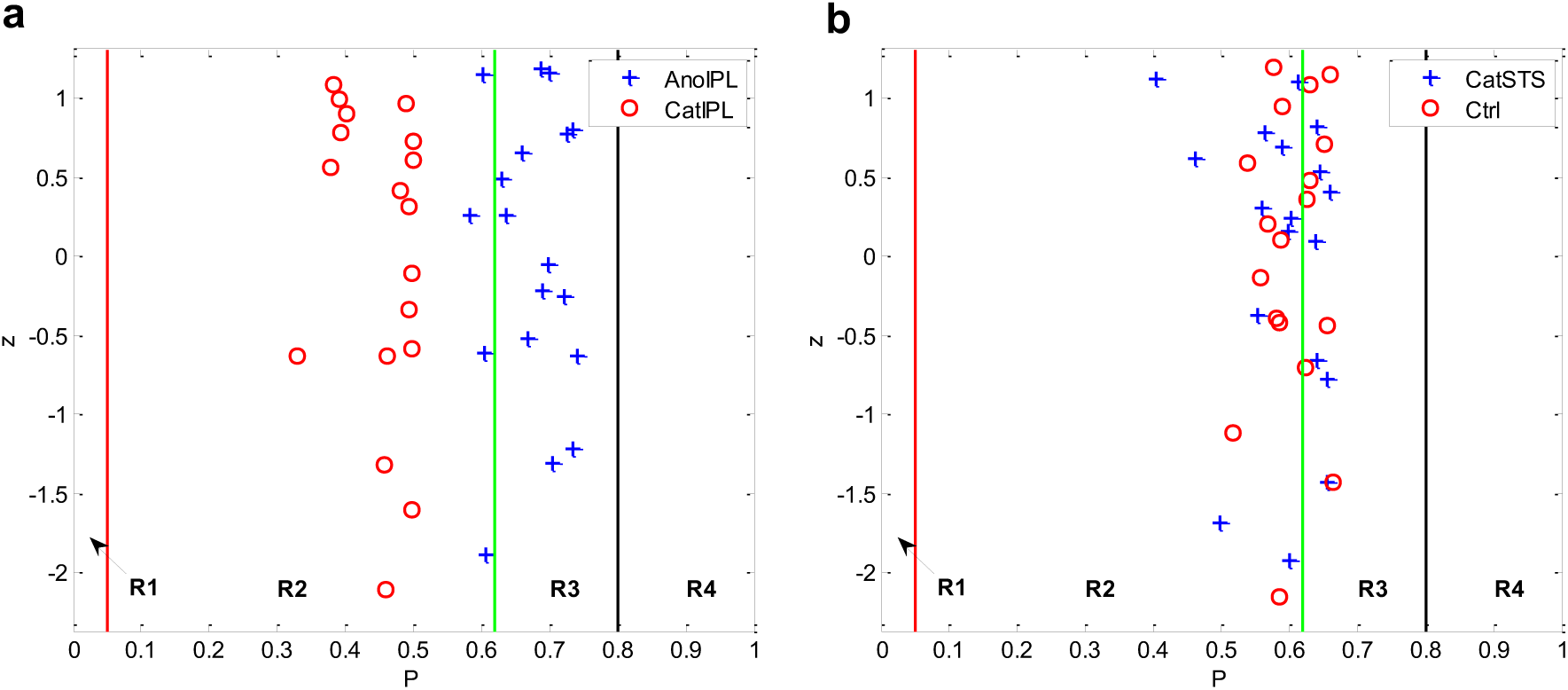
Plots of within-module degree *z* against participation coefficient *P.* For the four experimental groups (AnoIPL, CatIPL, CatSTS and Ctrl), *P*-*z*-plots are shown. The borders of the different regions (R1 – R4, see Methods) are marked by lines. There is a clear difference in distribution between groups AnoIPL and CatIPL (a) while there is no difference between groups CatSTS and Ctrl (b).

There is a clear difference visible in the distributions of *P* values between CatIPL and AnoIPL experimental groups (Figure 4 (a)) while there seems to be no difference in the other two experimental groups (Figure 4 (b)). Therefore, we analysed the differences in *P* distributions using the Wilcoxon signed-rank test as implemented in MATLAB R2013a. The one-tailed Wilcoxon signed-rank test with α = 0.01 shows a significant difference in AnoIPL > CatIPL (*z*_*wilcoxon*_ = 3.70, *p* << 0.01), AnoIPL > CatSTS (*z*_*wilcoxon*_ = 3.09, *p* << 0.01), AnoIPL > Ctrl (*z*_*wilcoxon*_ = 2.92, *p* << 0.01), CatIPL < CatSTS (*z*_*wilcoxon*_ = −3.66, *p* << 0.01) and CatIPL < Ctrl (*z*_*wilcoxon*_ = −3.70, *p* << 0.01). There was no significant difference using the two-tailed Wilcoxon signed-rank test with α = 0.01 in CatSTS ≠ Ctrl (*z*_*wilcoxon*_ = −0.85, *p* > 0.39).

Compared to both control experimental groups, more nodes in the AnoIPL experimental group jumped to region R3 while those of the CatIPL experimental group fell back completely to region R2. Finally, we analysed the differences in *z* distributions as well. There was no significant difference using the two-tailed Wilcoxon signed-rank test with α = 0.01 between groups: AnoIPL ≠ CatIPL (*z*_*wilcoxon*_ = 0.24, *p* > 0.81), AnoIPL ≠ CatSTS (*z*_*wilcoxon*_ = 0.20, *p* > 0.84), AnoIPL ≠ Ctrl (*z*_*wilcoxon*_ = −0.20, *p* > 0.84), CatIPL ≠ CatSTS (*z*_*wilcoxon*_ = 0.11, *p* > 0.91), CatIPL ≠ Ctrl (*z*_*wilcoxon*_ = 0.02, *p* > 0.98) and CatSTS ≠ Ctrl (*z*_*wilcoxon*_ = 0.37, p > 0.71). The role of nodes within their community (*z* value) does not differ significantly in all experimental groups. The role of nodes to other communities (P value) changed depending on the kind of stimulation. There was no change compared to Ctrl in the CatSTS experimental group. But in the AnoIPL experimental group, the community structure intensifies and so do the edges between communities. The four modules are more densely connected, the node roles jumped from region R2 (lower *P* values) to region R3 (higher *P* values) having more edges to other communities as compared to both control groups. The opposite is the case in the experimental group CatIPL where the module structure relaxes and so do the node roles. They drop completely to region R2 (lower *P* values) having less edges between communities than in both control experimental groups.

## Discussion

Here we show that tDCS to one node of a functional network affects the network architecture as a whole. The results presented here and in Almeida et al. (2017)5 provide a proof of principle that tDCS – delivered through the scalp using currents of 2 mA – can influence neuronal activity in humans. Moreover, they suggests that the effects of tDCS may arise from changed communication patterns (and not just local modulation of signal) that are modified by stimulation polarity and from altered functional connectivity between brain areas.

Crucially, our data shed light to some of the unresolved issues regarding the effects of tDCS at systems level. Namely, that: (i) tDCS is not limited to a local effect on the stimulated area, but exerts polarity-specific effects on the topology of the functional network attached; (ii) this effect is, if anything, only minimally affected by non-specific spread of the tDCS induced electrical field, but is rather dependent on network-specific processing of information; and (iii) at an intermediate scale, tDCS modulates functional connectivity by modular reorganisation.

Our results also show that in anodal tDCS the community structure in a regional and task-related network that is attached to the stimulation site intensifies and this leads to more edges between these communities. The existence of some edges between nodes in different communities acts as topological short-cuts38. This is in line with the results by Polania et al. (2011)23 who came to the conclusion that anodal tDCS increased the functional coupling between left somatomotor cortex (SM1) and neighboured topological regions (left premotor, motor and left parietal cortex) while the number of direct functional connections from left SM1 to topologically distant grey matter voxels decreased significantly. Interestingly, our results contradict Mancini et al.24 who stated that although tDCS is able to change network properties, it does not seem to affect the topological organisation of brain activity at a global level - which is not the case, as we show here.

Our results in the human brain are in line with those of Krause et al. (2017)20 who came to the conclusion that tDCS, in the primate brain, acts by modulating functional connectivity between brain areas. Despite the fact these authors showed – in agreement with Vöröslakos et al. (2018)13 – that in standard tDCS protocols, the electric field reaching the brain is too weak to alter the firing rate of neurons, they also detected a significant increase in anodal stimulation in the local field potential power and coherence in the targeted region when inspecting the effect of tDCS within the same protocols on the brain of living macaques – an ideal model system because of their thick, dense skull and gyrencephalic cortex similar to humans.

Finally, our data are highly consistent with the proposal that effects of tDCS depend on the level of ongoing activation in the particular functionally-defined target network47 – when we stimulated a node from another functionally-defined network (i.e., STS) we do not see any tDCS stimulation effects on the tool network.

To conclude, our findings confirm that tDCS influences neuronal activity in humans in a polarity-specific way, and does so in an experimental condition where participants are blind to the polarity of the tDCS stimulation, the measurement (BOLD signal) is bias free in what concerns the status of tDCS – i.e., within a completely independent analysis – and the neural tissue is alive and is engaged in processing incoming stimuli. Moreover, we also show that the flow of information within a functionally-isolated network is altered in a polarity-specific way and that this may be partially the locus of the causal relation between brain states and behaviour.

## Methods

### Data Acquisition and Pre-processing

We performed a combined tDCS/fMRI experiment on ten healthy adults at a 3T MAGNETOM Trio whole body MR scanner (Siemens Healthineers, Erlangen, Germany). The study adhered to the Declaration of Helsinki and was approved by the Ethic Committee of the Faculty of Medicine, University of Coimbra, Portugal. All participants gave written informed consent after a detailed description of the complete study. Participants went through four experimental sessions: a control session where they participated only in the fMRI experiment; a tDCS anodal session on IPL followed immediately by the fMRI experiment; a tDCS cathodal session on IPL followed immediately by the fMRI experiment; and a tDCS cathodal session on STS followed immediately by the fMRI experiment. All participants went through the control session first. The order of the tDCS sessions was counterbalanced across participants. Each session was separated by at least a week. During the fMRI experiment, the participants viewed pictures passively in an object processing paradigm (see 5) where we presented images of tools, animals, famous faces, and famous places in a miniblock design48 (each miniblock was restricted to a category). Within each run, miniblocks were pseudo-randomised; all participants completed five runs of this experiment which resulted in recording 455 functional volumes per session.

For analysis of functional brain networks, we extracted the overall mean time series from each of 18 brain regions known to be part of the tool network29,34,35,49,50,51,52 (see Table 1) using a BrainVoyager software (Brain Innovation, Maastricht, The Netherlands) adapted Anatomy Toolbox53. Before extraction of the time series, the functional volumes were pre-processed using BrainVoyager QX 2.8 applying slice-time and 3D motion correction, normalisation to Talairach space54, and z-normalisation. The time series were high-pass filtered (0.008 Hz) to remove low-frequency scanner drift before constructing functional brain networks.

### Construction of Functional Brain Networks

Each of the 18 ROIs selected above represents a single node in the resulting functional network. From the overall mean time series, we then obtained a temporal correlation matrix (size 18 × 18) for each participant by computing the Pearson partial correlation coefficients with controlled variables as implemented in MATLAB R2013a between time series of every pair of ROIs, while controlling for effects of noise. As covariates of non-interest for noise correction, we grouped the mean time series from white matter and cerebrospinal fluid extracted for each participant individually along with each participant’s motion parameters derived from the realignment step in pre-processing and the effects of the paradigm. The covariate of the paradigm effect was generated by convolving the box-car functions of paradigm conditions with the standard hemodynamic response function implemented in Statistical Parametric Mapping software (SPM12 (v6685), Wellcome Trust Centre for Neuroimaging, Institute of Neurology, University College London, UK) and was used to remove signal fluctuations of paradigm conditions from the time series. For each temporal correlation calculated, a p-value is given based on Student’s t distribution. To minimise the number of false-positives, we used a significance level of p < 0.002 (Bonferroni correction) to threshold the temporal correlation matrix of each participant. The remaining correlations can be interpreted as connections or edges between the nodes of the functional network. Here, the values of the correlation coefficients serve as edge weights showing the strength of a relation. While binary values enhance contrast they may also hide important information as edge weights below or above threshold may vary substantially between groups. Weighted graph analysis preserves this information. In our analyses, to avoid negative edge weights we converted them to absolute values because we were interested in any changes between the four experimental groups. It was shown elsewhere55 that linearly mapping the weight range [-1,1] to [0,1] kept the topology metrics of functional brain networks.

### Averaging correlation coefficients

There are at least three different methods to average correlation coefficients: (i) calculation of arithmetic mean of rs which is known to underestimate the true sample mean, (ii) Fisher’s z-transform and inverse Fisher’s z-transform before and after averaging which is known to overestimate the true sample mean56 and (iii) Olkin-Pratt estimator57 which is supposed to be least biased. Because of our sample sizes (N ≤ 10) which are known to be affected by bias58 most, we calculated averaged correlation matrices for each group using all three methods. Then, we computed all graph theory metrics listed below with the three group means averaged differently. There were no qualitative differences in the results. The choice of method had no noteworthy influence. For further analysis, we used the Olkin-Pratt estimator because it is recommended for averaging correlations either across samples or over repeated measures within sample59.

### Graph Theory Metrics

In general, networks (or graphs) are represented as sets of nodes *N* and edges *k*. Graphs are said to be unweighted if edges are either only present or absent – or weighted if edges are assigned weights. Graphs are undirected if edges do not contain directional information and directed if they do. Here, we analysed weighted undirected graphs by means of graph theory using the Brain Connectivity Toolbox41 (BCT, version 2017-01-15). All graphs analysed are connected graphs. Graph theory metrics depend on the number of *N* and k60 (*N,k*-dependence) as well as on the choice of correlation matrix and edge weights61. N,k-dependence can have two effects on graph theory metrics: (i) true effects are masked by opposite effects and (ii) significant effects are introduced. Here, we have primarily looked at graph theory metrics that are less sensitive to changes in *N* and *k* like topological metrics. First, we compared the graphs of the four groups concerning number of edges to address *N,k*-dependence of graph metrics. The number of nodes (here: 18) stays constant throughout groups. Then, we looked at topological metrics such as modularity, community structure, within-module degree *z* score, participation coefficient and distance.

#### Degree

Node degree is the number of links connected to a node. During calculation of node degree using BCT, weight information on edges is discarded60.

#### Modularity

The modularity *Q* measures the goodness with which a graph is optimally partitioned into functional subgroups or communities. For weighted graphs, modularity is defined as62

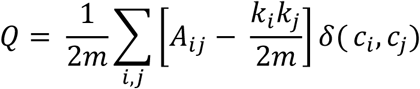

with *A*_*ij*_: weight of edge between *i* and *j, k*_*i*_ = ∑_j_ *A*_*ij*_: sum of weights of edges attached to vertex *i, c_i_*: community vertex *i* is assigned to, δ(*x,y*) is 1 if *x* = *y* and 0 otherwise and *m* = 1/2 ∑_ij_ *A*_*ij*_. Being a scalar value, *Q* lies in the interval [‒1,1], theoretically. If the fraction of within-community edges is no different from what is expected for the randomised network, then *Q* will be zero. Nonzero values indicate deviations from randomness. *Q* measures the density of links inside communities compared with links between communities. In this context, the modularity *Q* is used as an objective function to optimise during graph partitioning: the higher the value of *Q* the better the partitioning. If the number of edges within communities exceeds the number of edges expected by chance the value of *Q* is positive.

#### Community structure

If nodes of a graph can be easily partitioned into sub-units of densely connected nodes, the graph is presumed to have community structure. This implies that communities merely consist of nodes with more densely connections within and more sparsely connections between communities. This definition only holds true for positive edge weights in the first place. Concerning negative edge weights, the assignment of nodes should be done the opposite way compared to positive edge weights, that is negative edges are sparse within and more dense between communities45, a concept evolving from social balance theory63. Although we computed all graph theory metrics using absolute values we cross-checked this limitation by overlaying the community structure for each group on their averaged weighted temporal correlation matrix before converting it to absolute values to verify this issue. As specified before, modularity is an objective function measuring the quality of a graph’s community partition. By searching over all possible partitions of a graph, the modularity optimisation method identifies communities that have a high modularity value *Q*. The detection of a graph’s optimal community structure is essential as it may identify functional sub-units so far unknown that influence the overall behaviour of the graph. The optimal community structure is a partition of the graph into non-overlapping sub-units of nodes maximising the number of edges within sub-units and minimising the number of edges between sub-units44. One limitation of modularity optimisation is the resolution limit64 which could lead to failure in resolving even well-defined small communities. Therefore, it might be possible that communities found are clusters of communities in fact. This might be the case if 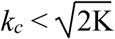 where *k*_*c*_ denotes the number of internal edges in the community *c* and *K* the total number of edges in the graph. Therefore, it is important to look more closely at the internal structure of all communities found as can be done by using the inequation44

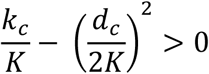

with *d*_*c*_: total degree of nodes in community. If the inequation holds true the community under consideration is actually a single community and not a mixture of two or more smaller ones. All communities found in our analysis comply with the inequation given above. Because community detection using exact modularity optimisation is an NP-hard problem, BCT implemented the Louvain algorithm64 which contains a stochastic element that lets the output vary from run to run. To account for this issue, we ran the algorithm a 1000 times per group and used consensus clustering65 for selection of best community structure for further computations. Once the community structure of a graph is known, the following two graph theory metrics are easily computed.

#### Within-module degree z score

The internal organisation of a community or module may vary between totally centralised nodes (one or a few nodes connected to all the others) and totally decentralised ones (all nodes having similar number of edges). Nodes are said to fulfil similar roles if they have similar connectivity within a community. The within-module degree z-score is a metric of how well-connected a node is to other nodes in a community46 and is defined as

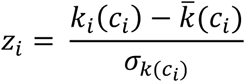

with *c*_*i*_:module containing node *i, k_i_*(*c*_*i*_): within-module degree of 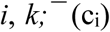: mean of within-module *c*_*i*_ degree distribution and σ_k(*ci*)_: standard deviation of the within-module *c*_*i*_ degree distribution. The higher the values of *z*, the higher the within-module degrees are and vice versa which implies that nodes with *z* ≥ 2.5 can be classified as hub nodes and nodes with *z* < 2.5 as non-hub nodes46. Both types of nodes can be subdivided even further by using the values of the participation coefficient P.

#### Participation coefficient

The two areas in the *z*-plane (hub and non-hub nodes) can be fine-grained because of the connections of a node to communities other than its own. Sharing the same *z*-score, one node might be connected to several nodes in other communities while the other might not. The participation coefficient acts as a measure of diversity of inter-modular connections of nodes46 and is defined as

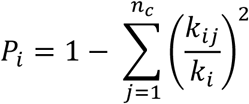

with *k*_*ij*_: number of edges of node *i* to nodes in community *j, k_i_*: total degree of node *i* and *n*_*c*_: number of communities detected. The participation coefficient *P* measures how ‘well-distributed’ the edges of a node are among different communities. It is close to 1 if the edges are uniformly distributed among all the communities and 0 if the entire edges are within its own community.

#### Node topology

Based on the idea that nodes with the same role should have similar topological properties, the role of a node can be determined by its location in the *P-z-*parameter plane which defines how the node is positioned in its own community and relative to others. Guimerà and Amaral46 defined seven regions by dividing the *P–z* parameter plane in different areas. Because we are only looking at the tool network, we do not expect to find any hub nodes (*z* ≥ 2.5). So, here we only took into account the non-hub nodes area (*z* < 2.5) that was subdivided into four different regions: R1 – nodes with all their edges within their module (*P* ≤ 0.05); R2 – nodes with at least 60% of their edges within their module (0.05 < *P* ≤ 0.62); R3 – nodes with half of their edges to other modules (0.62 < *P* ≤ 0.80); and R4 – nodes with edges homogeneously distributed among all modules (*P* > 0.80). Such nodes were classified as kinless nodes and are said to be mostly found in network growth models, but not in real-world networks.

#### Distance

The distance matrix shows the length of shortest paths between all pairs of nodes. Each entry stands for the number of edges that have to be traversed to get from one node to another. By using a weighted correlation matrix, higher correlation coefficients denote shorter distances. We converted the weighted correlation matrices to length by inversion of weights and fed them into Dijkstra algorithm66 to compute the distance between nodes.

## Acknowledgements

This research was supported by a Foundation for Science and Technology of Portugal and Programa COMPETE grant (PTDC/ MHC-PCN/0522/2014) to J.A. M.R. and J.A. were supported by Deutscher Akademischer Austauschdienst (Projekt-ID 57212180) and CRUP. S.K. is supported by a Foundation for Science and Technology of Portugal and Programa COMPETE grant (PTDC/MHC-PCN/0522/2014). This research was supported by German Research Foundation grant SCHA 546/22-1 to L.R.S.

## Author contributions

M.R. performed data analyses, prepared the graphics and wrote the paper. S.K. performed data analysis. L.R.S. contributed ideas. J.A. conceived and designed the project, and critically revised the manuscript. M.R. and J.A. interpreted the data.

